# Molecular Crowding Suppresses Mechanical Stress-Driven DNA Strand Separation

**DOI:** 10.1101/2024.12.11.628023

**Authors:** Parth Rakesh Desai, John F. Marko

## Abstract

Molecular crowding influences DNA mechanics and DNA - protein interactions and is ubiquitous in living cells. Quantifying the effects of molecular crowding on DNA supercoiling is essential to relating *in-vitro* experiments to *in-vivo* DNA supercoiling. We use single molecule magnetic tweezers to study DNA supercoiling in the presence of dehydrating or crowding co-solutes. To study DNA supercoiling, we apply a stretching force of 0.8 pN to the DNA and then rotate one end of the DNA to induce supercoiling. In a 200 mM NaCl buffer without co-solutes, negatively supercoiled DNA absorbs some of the tortional stress by forming locally melted DNA regions. The base-pairs in these locally melted regions are believed to adopt a configuration where nucleotide base pairing is disrupted. We find that the presence of a dehydrating co-solute like glycerol further destabilizes base-pairs in negatively supercoiled DNA. The presence of polyethylene glycol, commonly used as a crowding agent, suppresses local strand separation and results in plectoneme formation even when DNA is negatively supercoiled. The results presented in this letter suggest further directions for studies of DNA supercoiling and supercoiled DNA – protein interactions in molecular conditions that approximate *in-vivo* molecular composition.

**SIGNIFICANCE:** Accurate modelling of DNA mechanics is central to interpreting results of single molecule studies of DNA mechanics and DNA-protein interactions. While the effect of molecular conditions on thermal stability of short and relaxed DNA has been studied, the influence of molecular conditions on DNA supercoiling has not been explored. We present the first single molecule study of DNA supercoiling in the presence of crowding and dehydrating co-solutes. We observe that co-solutes can increase or completely suppress mechanical stress-driven base-pair disruption in negatively supercoiled DNA. This change of DNA supercoiling is likely to significantly affect the function of DNA-binding proteins. Our results motivate the need for systematic exploration of DNA supercoiling in presence of co-solutes to accurately relate *in-vitro* DNA-protein interactions to *in-vivo* DNA-protein interactions.

## INTRODUCTION

*In-vivo*, molecular conditions surrounding DNA are varied. In *Escherichia coli* (*E. coli*) cell, DNA is negatively supercoiled and surrounded by 30-40% volume fraction of other organic and inorganic solutes(1, 2). The presence of protein complexes also changes the molecular conditions around DNA. Previous studies have demonstrated that co-solutes affect thermal stability of small and relaxed DNA molecules(3, 4). Previous studies have shown that molecules that bind to DNA can change the supercoiling behavior of DNA(5, 6). However, the effect of non-binding co-solutes on DNA supercoiling has not been systematically studied.

The presence of glycerol and ethylene glycol has been demonstrated to reduce the melting temperature of small DNA molecules(7–9). Previous studies have observed that glycerol and ethylene glycol reduce the thermal stability of short DNA molecules because they replace some of the water molecules surrounding the DNA(7, 10, 11). In addition to thermal destabilization, presence of co-solutes can also impact specificity of DNA-protein interactions(12–14).

In the presence of crowding agents like polyethylene glycol (PEG), depending on the size of both PEG and DNA both thermal stabilization and thermal destabilization have been observed(15). Single-molecule magnetic tweezers experiments have also previously observed that DNA can be osmotically compacted by PEG molecules(16–19). The amount of compaction is affected by the counterion valency and the size of PEG molecule(16). Studies of PEG-DNA interactions have also suggested presence of PEG causes a decrease in energy required to form plectonemes(20). PEG molecules can also induce local melting in DNA molecules(21).

In this letter, we investigate the effect of dehydrating and crowding co-solutes on DNA supercoiling. Using single molecule magnetic tweezers(22), we observe that presence of co-solutes can differently affect DNA’s response to thermal and mechanical stress. As for studies of thermal melting, the presence of glycerol increases local strand separation in mechanically stressed DNA. The presence of ethylene glycol did not significantly affect DNA supercoiling, while presence of its polymer. polyethylene glycol (PEG) 8000 suppresses base pair separation in negatively supercoiled DNA. We also use the extension-rotation curves to quantitatively calculate the mechanical parameters required to model DNA supercoiling in presence of different co-solutes. The results presented here demonstrate that *in-vivo* DNA supercoiling is influenced by molecular conditions by a combination of chemical and physical effects. Since molecular conditions in living cells are highly crowded, and DNA-protein interactions are sensitive to DNA structure, accurate extrapolation of *in-vitro* DNA-protein interactions studies to *in-vivo* phenomenon requires consideration of the influence of co-solutes on DNA supercoiling.

## RESULTS AND DISCUSSION

To study the effects of co-solutes on DNA supercoiling, we use single molecule magnetic tweezers and measure the change in extension of DNA due to supercoiling in presence of different co-solutes. We start by measuring the extension-rotation curve in a buffer of 200 mM NaCl. We then flow in a solution of 200 mM NaCl plus specific co-solutes (see additional details in supplementary section S1). The extension-rotation curves are measured with a ~10 kb DNA under a force of 0.8 pN. In the salt concentration and force condition used here, the extension-rotation curve is asymmetric (blue curves in Fig. 1). For DNA overtwisting (positive supercoiling), DNA absorbs the excess turns by increasing its internal twist, until a critical number of turns is absorbed. Once one passes that critical number of turns, subsequent turns cause the DNA to buckle and form a plectonemic loop(23, 24). Each subsequent turn causes elongation of the plectoneme and results in a corresponding reduction in DNA extension. In the under-twisting (negative supercoiling) regime, under the force and salt conditions used here, DNA absorbs the applied turns through a combination of local strand separation and plectoneme formation(25).

**FIG 1.**
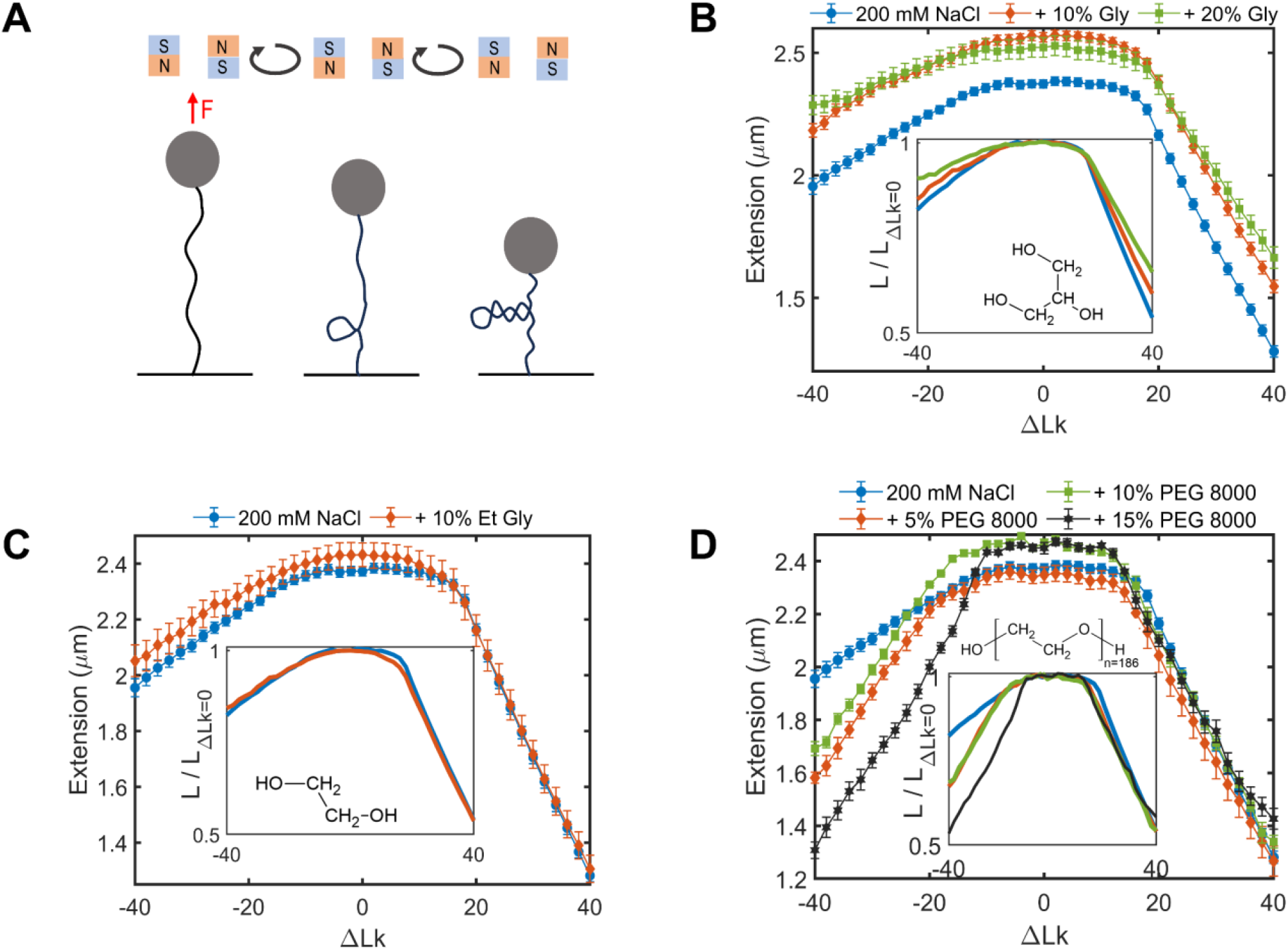
Co-solutes affect DNA supercoiling. (A) Schematic of MT setup. A paramagnetic bead attached to the DNA molecule is used to apply stretching force to the DNA molecule. Buckling can be induced by rotating the paramagnetic bead. (B) Extension-rotation curve of DNA in 200 mM NaCl and 0% (blue), 10% (red) and 20% (green) v/v Glycerol (Gly) buffer. (C) Extension-rotation curve of DNA in 200 mM NaCl and 0% (blue) and 10% (red) v/v Ethylene Glycol (Et Gly) buffer. (D) Extension-rotation curve of DNA in 200 mM NaCl and 0% (blue), 5% (red), 10% (green) and 15% (black) wt./v Polyethylene Glycol 8000 (PEG) buffer. Insets show chemical structure of co-solute, and the extension-rotation curve normalized to the extension of a tortional relaxed DNA in the respective buffer conditions. Error bars represent standard errors. Chemical structures were created using ChemSketch(27).

Marko *et al*. (23) models DNA supercoiling as phase coexistence between stretched DNA and plectonemic DNA and provides closed-form equations to determine mechanical parameters from the extension-rotation curve. Quantitative modeling of DNA supercoiling can be performed using the bending persistence length of relaxed DNA (*A*), twist persistence length of DNA (*C*) and plectoneme-linking number stiffness (*P*, termed twist stiffness of plectoneme in Ref. *(23)*). We determine *C* and *P* from the critical buckling linking number and slope of the extension-rotation curve (see supplementary section S4). We generally obtain *C* and *P* from extension-rotation curve of positively supercoiled DNA. Critical buckling linking number is not well defined for negatively supercoiled DNA in the force and salt condition used here. An important exception is negatively supercoiled DNA in presence of PEG 8000, where we can observe a critical buckling linking number for negatively supercoiled DNA. The values of *C* obtained from the method described here, can be physically interpreted as an indicator of the twist rigidity at the critical buckling linking number. For positively supercoiled DNA, *C* and *P* represent properties intrinsic to the double helix, while for negatively supercoiled DNA, *C* can be reduced by base-pair destabilization(23). The *C* (~85 nm) and *P* (~20 nm) values of positively supercoiled DNA in 200 mM NaCl buffer are comparable to recent direct measurements by Gao et *al*. (26). These parameters can be used to create quantitative models of DNA supercoiling in the presence of different co-solutes.

In the presence of a dehydrating co-solute like glycerol, we observe a decrease in slopes of the extension-rotation curves for both positively and negatively supercoiled DNA (see Fig. 2 (A)). The change in slope can be interpreted as a reduction in *C* and *P* (see Fig. 2 (C) and 2 (D)), suggesting that some of the applied linking number are absorbed in twist which results in a shorter plectoneme and hence a more extended supercoiled DNA. Previous studies of DNA response to thermal stress have shown that presence of glycerol reduces the melting temperature of short and relaxed DNA(7–9). Our results indicate a similar destabilization effect of glycerol for DNA under mechanical stress. As in prior experiments, we also observe an increase in DNA extension of relaxed DNA due to glycerol(28). Our results suggest that glycerol alters the base-pair structure of relaxed and supercoiled DNA. Glycerol is a common additive utilized in *in-vitro* experiments of DNA-protein interactions. The change in DNA supercoiling due to glycerol can affect energetics of DNA-protein interactions.

**FIG 2.**
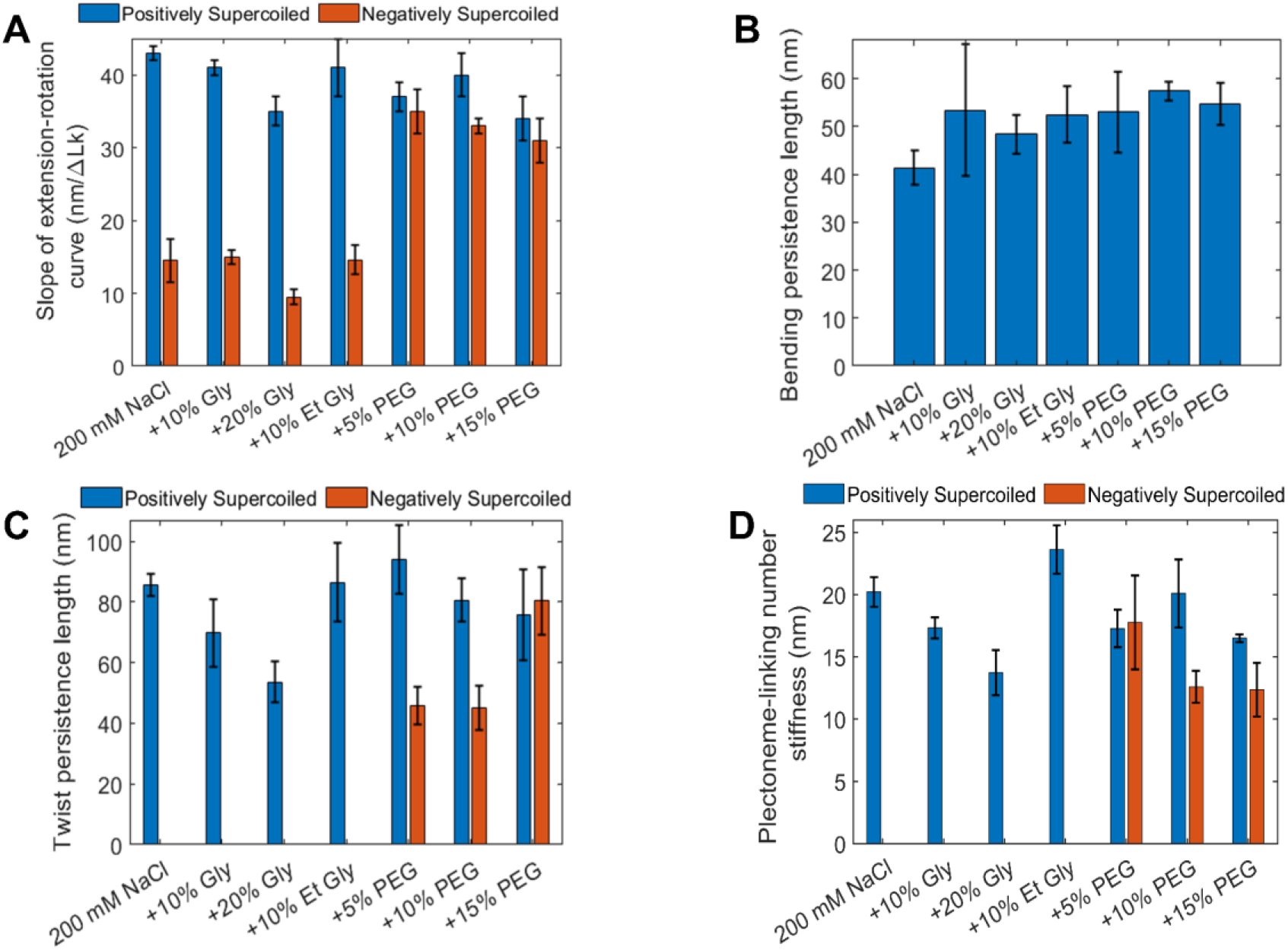
Statistical-mechanical parameters associated with DNA supercoiling. (A) Average slope of extension-rotation curves of positively supercoiled (blue) and negatively supercoiled DNA (red) in presence of co-solutes. (B) Average bending persistence length of DNA in presence of co-solutes. (C) Average twist persistence length of DNA measured from extension-rotation curves of positively supercoiled (blue) and negatively supercoiled (red) DNA. (D) Average plectoneme-linking number stiffness measured from extension-rotation curves of positively supercoiled (blue) and negatively supercoiled (red) DNA. Error bars represent standard errors from independent measurements.

To test if DNA local strand separation due to mechanical stress can be exacerbated by other de-hydrating agents, we measured extension-rotation curves in presence of ethylene glycol. Previous studies have observed that presence of ethylene glycol decreases the melting temperature of short DNA molecules(8). In single molecule magnetic tweezers experiments, we do not observe significant change in slope of the straight portions of the extension-rotation curve, *C* or *P* (see Fig. 1 (C) and Fig. 2). Our results indicate that thermal destabilization may not always indicate a corresponding mechanical destabilization.

Next, we investigate the effect of polymerized ethylene glycol, polyethylene glycol (PEG), on DNA supercoiling. Previous studies of small DNA molecules have observed that lower molecular weight PEG molecules reduce the melting temperature, while higher molecular weight PEG molecules increase the melting temperature of small DNA(15). Driven by osmotic exclusion, PEG molecules have been observed to induce compaction and condensation in long DNA molecules(16). Higher molecular weight PEG molecules can induce compaction at lower PEG concentrations. Single molecule magnetic tweezers have previously observed salt dependent compaction of relaxed DNA in the presence of PEG(16). The presence of a crowding agent has been shown to enhance the activity of DNA binding proteins. These studies indicate that crowding can play an important role in protein activity(19, 29); however, the effect of crowding on supercoiled DNA has not been studied.

Using single molecule magnetic tweezers, we observe that the presence of PEG 8000 suppresses local strand separation in negatively supercoiled DNA. The asymmetry in slopes on the two sides of the extension-rotation curve can be used to estimate the suppression of local strand separation in supercoiled DNA due to PEG 8000. Adding 5%, 10% or 15% wt./v PEG 8000 to 200 mM NaCl buffer decreases the asymmetry in slopes of extension-rotation curve, suggesting that helix opening is suppressed in these conditions (see Fig. 2 (A)). The presence of a well-defined critical buckling linking number for negatively supercoiled DNA in presence of PEG 8000, allows us to calculate *C* and *P* from the extension-rotation curve. We observe that in presence of 15% wt./v PEG 8000, *C* is similar for both positively and negatively supercoiled DNA, suggesting that twist rigidity is restored in crowded conditions. In presence of PEG 8000 we also observe a decrease in *P*, suggesting that molecular crowding decreases energy required to elongate plectonemic region in DNA. We also find that addition of PEG results in plectoneme formation at a lower linking number. Generally, plectoneme formation due to torsional stress can be modelled as a competition between twisting and bending(23). We observe only a moderate increase in persistence length of relaxed DNA in the presence of PEG 8000 (see Fig. 2 (A)), suggesting that the decrease in critical buckling linking number is driven by change in *C* and *P*.

Given the relative size of PEG 8000 (~4.5 nm(30, 31)) and plectoneme loop (~40 nm), one might be tempted to attribute the shorter extension of negatively supercoiled DNA in PEG buffer to compression of plectoneme structure caused by osmotic exclusion. However, if osmotic exclusion was a significant factor in the decrease in extension, we ought to have also observed a similar effect of PEG on positively supercoiled DNA. The fact that these effects are absent for similar concentrations of ethylene glycol are consistent with a physical molecular-crowding-based mechanism rather than a chemical interaction one.

## CONCLUSION

We present the first single molecule analysis of DNA supercoiling in presence of a high concentration of co-solutes. We observe that thermal destabilization of DNA in presence of co-solutes does not necessarily evidence mechanical destabilization of DNA in similar conditions. We also observed that local strand separation in negatively supercoiled DNA can be suppressed due to presence of crowding co-solutes like PEG 8000. Our results indicate that relating *in-vitro* studies of DNA-protein interactions to *in-vivo* phenomena requires consideration of the change in DNA mechanics in presence of co-solutes; for example, changes to the mechanical properties of DNA (bending and twisting persistence lengths) will result in different behaviors for enzymes which act to bend, twist, or change linking number of the double helix.

## AUTHOR CONTRIBUTIONS

J.F.M. supervised the project. J.F.M. and P.R.D. designed the project. P.R.D. performed experiments and analyzed the data. P.R.D. wrote the manuscript. J.F.M. and P.R.D. read, revised and approved the manuscript.

## ACKNOWLEDGEMENT

This work was supported by NIH grant R01-GM105847, and by subcontract to the University of Massachusetts under NIH grant UM1-HG011536 (Center for 3D Structure and Physics of the Genome, 4DN Consortium.

## Supporting Information for

### S1. Materials and methods

#### DNA Preparation

The DNA substrate was constructed by ligating functionalized digoxigenin and biotin handles (~ 900 bp long) to either end of a linearized pFOS1 plasmid (~ 10 kb). Modified DNA molecules were incubated with the M270 streptavidin-coated magnetic beads. DNA molecules were then incubated in a ~ 40 uL volume flow cell containing anti-digoxigenin functionalized surface. A more detailed protocol of DNA and flow cell construction can be found in Zaichuk et *al*. (1).

#### Procedure

The flow cell was then washed with > 10* flow cell volume of the standard salt buffer (200 mM NaCl, 20 mM TRIS-pH 7.5, 0.01% Tween 20). Supercoilable DNA molecules were found and verified by negatively supercoiling the DNA molecule under ~1 pN force and then reducing the force to ~0.2 pN. A rapid reduction in DNA extension would be observed if a single supercoilable DNA molecule was attached to the magnetic bead under observation. For the extension-rotation curves, we collected the DNA extension data at a rate of ~70 Hz for 20 seconds at every 2 turns between −40 and +40 turns.

Similar to previously published work(1), we calibrate the force applied to the DNA molecule by measuring the variation in horizontal motion of the magnetic bead for different magnet positions. We always start by measuring the extension-rotation curve for each DNA molecule in a 200 mM NaCl buffer. Next, the flow cells are washed with > 10* flow cell volume of the applicable co-solute buffer. After flowing in the new buffer, we always recalibrate the instrument to account for the change in refractive index due to presence of co-solute.

**Table S1.**
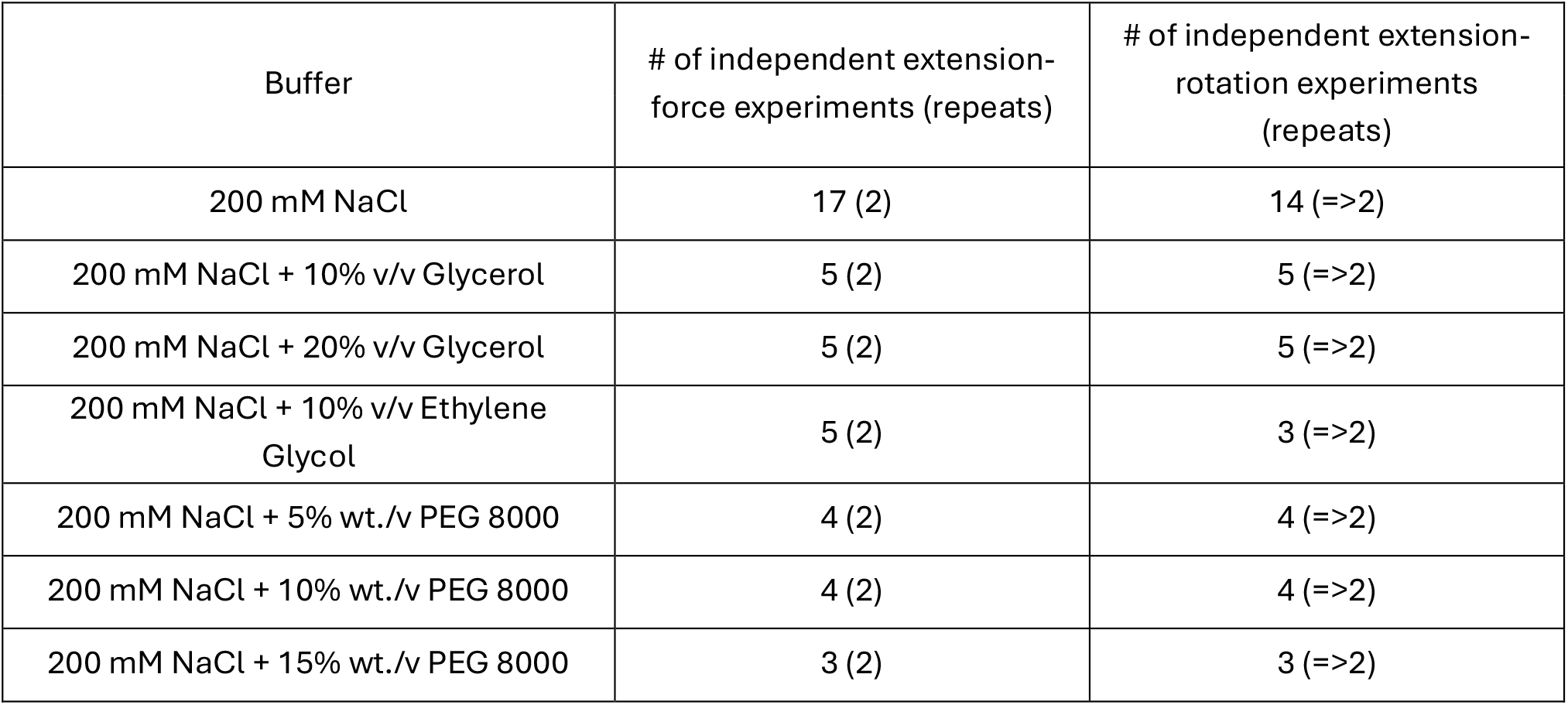
Details of independent experiments and repeats.

**Table S2.**
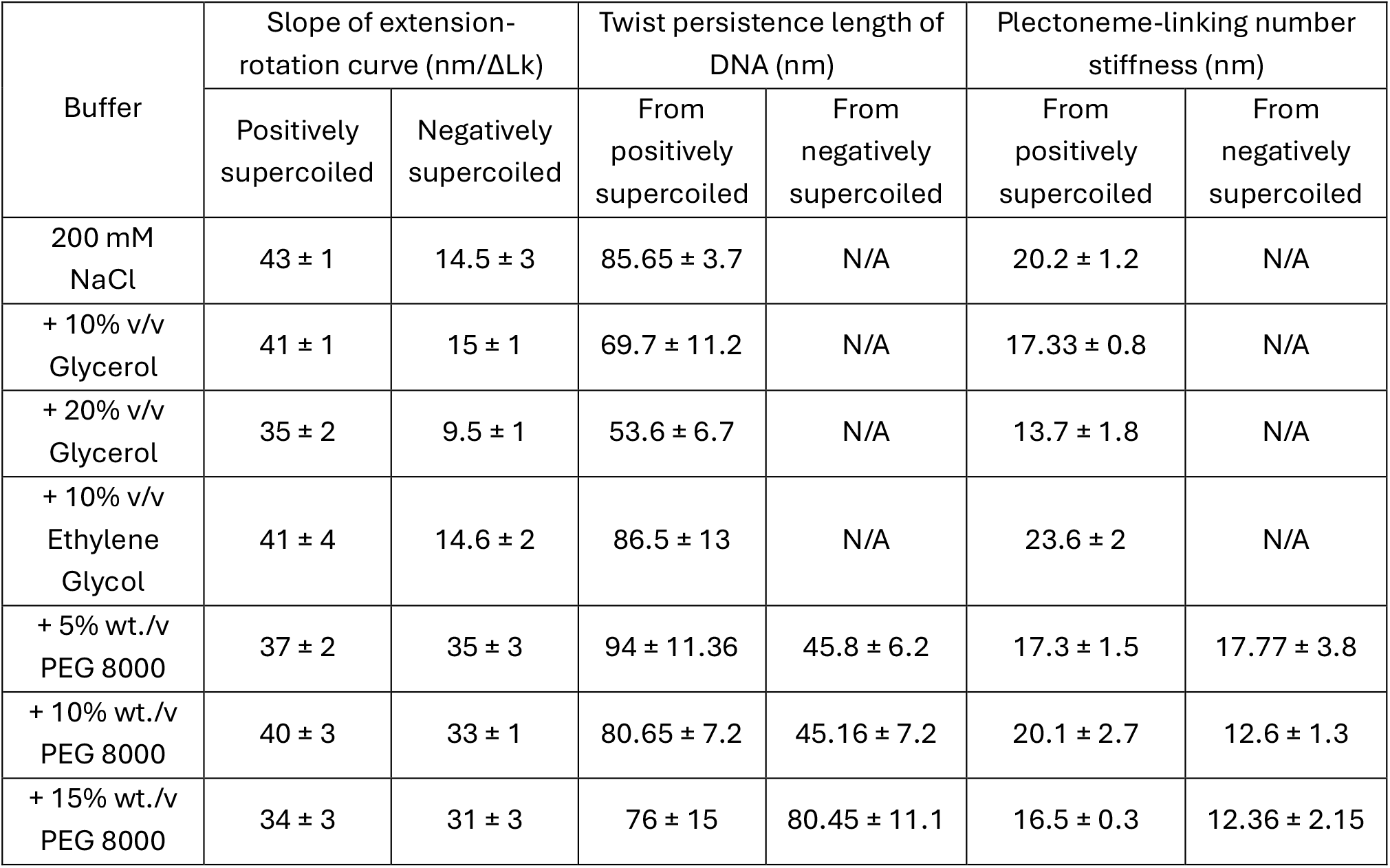
Calculated mechanical parameters required to model DNA supercoiling. Errors represent standard errors.

| Buffer | Slope of extension-rotation curve (nm/ΔLk) | | Twist persistence length of DNA (nm) | | Plectoneme-linking number stiffness (nm) | |
| --- | --- | --- | --- | --- | --- | --- |
|  | Positively supercoiled | Negatively supercoiled | From positively supercoiled | From negatively supercoiled | From positively supercoiled | From negatively supercoiled |
| 200 mM NaCl | 43 ± 1 | 14.5 ± 3 | 85.65 ± 3.7 | N/A | 20.2 ± 1.2 | N/A |
| + 10% v/v Glycerol | 41 ± 1 | 15 ± 1 | 69.7 ± 11.2 | N/A | 17.33 ± 0.8 | N/A |
| + 20% v/v Glycerol | 35 ± 2 | 9.5 ± 1 | 53.6 ± 6.7 | N/A | 13.7 ± 1.8 | N/A |
| + 10% v/v Ethylene Glycol | 41 ± 4 | 14.6 ± 2 | 86.5 ± 13 | N/A | 23.6 ± 2 | N/A |
| + 5% wt./v PEG 8000 | 37 ± 2 | 35 ± 3 | 94 ± 11.36 | 45.8 ± 6.2 | 17.3 ± 1.5 | 17.77 ± 3.8 |
| + 10% wt./v PEG 8000 | 40 ± 3 | 33 ± 1 | 80.65 ± 7.2 | 45.16 ± 7.2 | 20.1 ± 2.7 | 12.6 ± 1.3 |
| + 15% wt./v PEG 8000 | 34 ± 3 | 31 ± 3 | 76 ± 15 | 80.45 ± 11.1 | 16.5 ± 0.3 | 12.36 ± 2.15 |

### S2. Representative hat curves in presence of co-solutes

**FIG S1.**
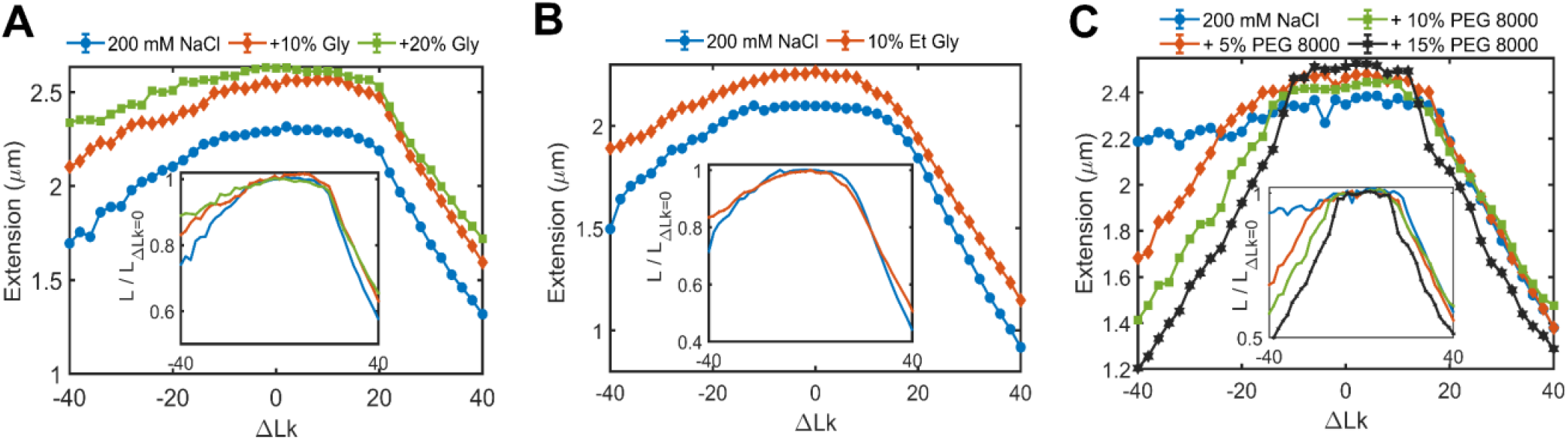
Representative extension-rotation curves demonstrate that presence of co-solutes affect DNA supercoiling. The extension-rotation curves presented in each sub-plot were collected on the same molecule. (A) Extension-rotation curve of DNA in 200 mM NaCl and 0% (blue), 10% (red) and 20% (green) v/v Glycerol (Gly) buffer. (B) Extension-rotation curve of DNA in 200 mM NaCl and 0% (blue) and 10% (red) v/v Ethylene Glycol (Et Gly) buffer. (C) Extension-rotation curve of DNA in 200 mM NaCl and 0% (blue), 5% (red), 10% (green) and 15% (black) wt./v Polyethylene Glycol 8000 (PEG) buffer. Inset show the extension-rotation curve normalized to the extension of a tortional relaxed DNA in the respective buffer conditions. Error bars represent standard errors and are smaller than the symbol size.

### S3. DNA bending persistence length and contour length in presence of co-solutes

**Figure S2.**
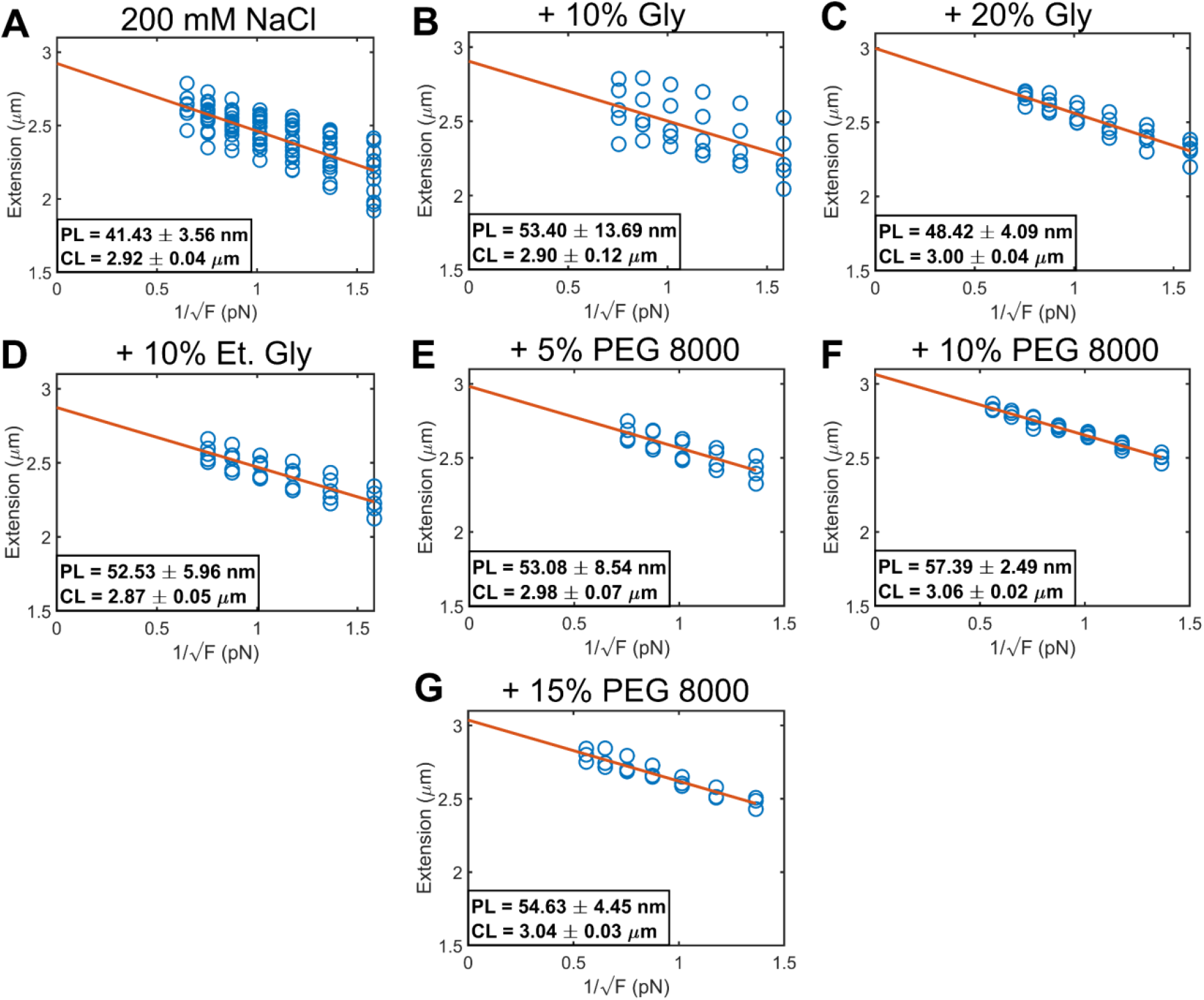
Persistence length (PL) and contour length (CL) of dsDNA in presence of different co-solutes. Persistence length was measured using the slope of the Extension-inverse of the square root of force curve(1). Extension (blue circles) was measure with measured at forces between ~0.8 pN and ~1.5 pN. Slope and intercept of the best linear fit line (red solid line) in plots of the DNA extension versus the inverse of the square root of force were used to determine the persistence length (PL), contour length (CL) and standard errors in accord with expectations from the worm-like chain polymer model.

### S4. Twist persistence length of DNA and twist stiffness of plectoneme

**Figure S3.**
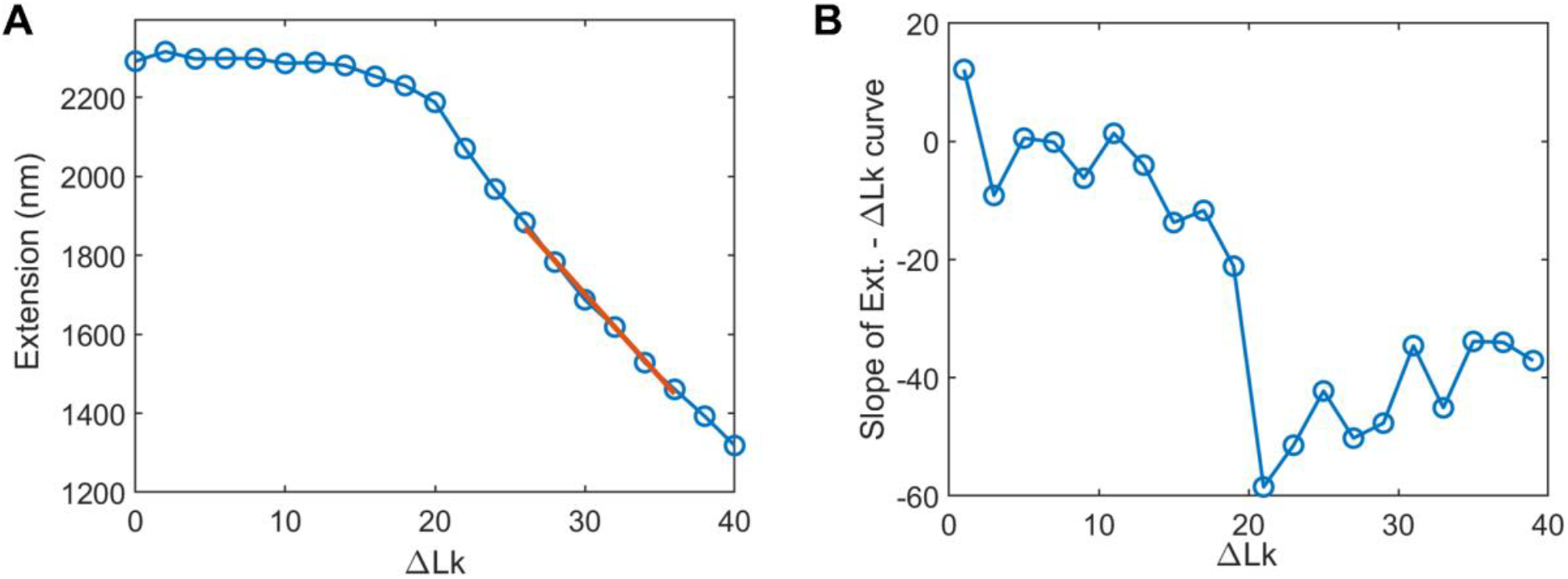
Extension-rotation curve is utilized to calculate the twist persistence length of DNA and plectoneme-linking number stiffness. (A) Representative extension-rotation curve of DNA under 0.8 pN of stretching force in 200 mM NaCl buffer. A straight line is fit to extension between ΔLk = 24 and ΔLk = 34. The line fit can be extrapolated to calculate the supercoiling density (σ_p_) required for zero extension. (B) Slope of the extension-rotation curve. Supercoiling density at buckling (σ_s_) can be calculated as minimum slope (first derivative) of the extension-rotation curve.

Quantitative modelling of DNA supercoiling can be performed using the bending persistence length of relaxed DNA (*A*), twist persistence length of DNA (*C*) and plectoneme-linking number stiffness (*P*) (plectoneme-linking number stiffness is similar to the “twist stiffness of plectoneme” parameter defined in Marko et *al*.(2)). The bending persistence length of DNA is calculated as described in supplementary section S3. Model presented in Marko et *al*.(2) can be used to obtain the twist persistence length of DNA and twist stiffness of plectoneme from the extension-rotation curve. Marko et *al*.(2) describes the supercoiling density at buckling (|σ_s_|) and supercoiling density for zero extension (|σ_*p*_|) as

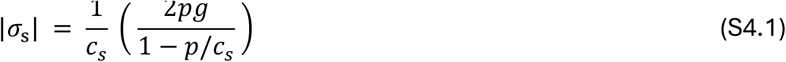

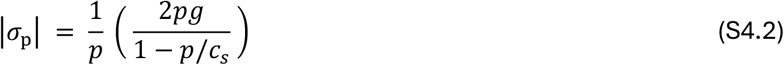

where, free energy of stretched and nicked DNA (*g*) under a stretching force (*f*) is 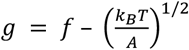. Twist stiffness of plectoneme 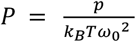 *&* the contour-length rate of rotation of the relaxed double helix 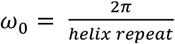.

The twist persistence length of DNA (*C*) is related to *c*_s_ and defined as 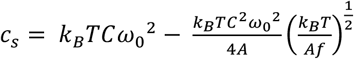. We obtain the buckling point (|*σ*_s_|) by finding minimum of the first derivative of the extension-rotation curve (see supplementary Fig. S3 (B)). The model defines |σ_*p*_| as the supercoiling density when all of the DNA molecule is part of plectoneme and hence DNA extension is zero. To calculate |σ_*p*_|, we first fit a straight line to the extension between ΔLk = 24 and ΔLk = 34. We then extrapolate the line to zero extension and calculate |σ_*p*_|.

After calculating |*σ*_s_| and |*σ*_p_| from the extension-rotation curve, we can divide Eq. S4.1 by Eq. S4.2 to get

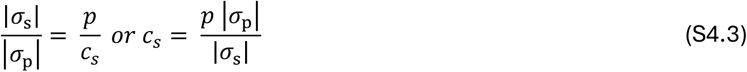

Eq. S4.3 can be used to simplify Eq. S4.1 to

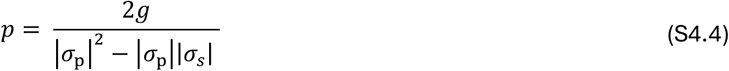

Eq. S4.4 and Eq. S4.3 can be used to obtain *p* and *c*_*s*_, which can then be used to twist persistence length (*C*) and plectoneme-linking number stiffness (*P*). The twist persistence length (~85 nm) and plectoneme-linking number stiffness (~20 nm) values of supercoiled DNA in 200 mM NaCl buffer are comparable to recent direct measurements by Gao et *al*. (3)

We only use |*σ*_s_| and |*σ*_p_| obtained from extension-rotation curve of positively supercoiled DNA because |*σ*_s_| is not well defined for negatively supercoiled DNA in the salt and force conditions used in the current paper.

