## Supplementary Information for "Molecular Crowding Suppresses Mechanical Stress-Driven DNA Strand Separation"

Parth Rakesh Desai^1^ and John F. Marko^1,2*^

^1^ Department of Molecular Biosciences, Northwestern University, Evanston, Illinois 60208, USA

^2^ Department of Physics and Astronomy, Northwestern University,
Evanston, Illinois 60208, USA

*****

**S1. Materials and methods**

**DNA Preparation**: The DNA substrate was constructed by ligating functionalized digoxigenin and biotin handles (~ 900 bp long) to either end of a linearized pFOS1 plasmid (~ 10 kb). Modified DNA molecules were incubated with the M270 streptavidin-coated magnetic beads. DNA molecules were then incubated in a ~ 40 uL volume flow cell containing anti-digoxigenin functionalized surface. A more detailed protocol of DNA and flow cell construction can be found in Zaichuk et *al*. (1).

**Procedure**: The flow cell was then washed with > 10* flow cell volume of the standard salt buffer (200 mM NaCl, 20 mM TRIS- pH 7.5, 0.01% Tween 20). Supercoilable DNA molecules were found and verified by negatively supercoiling the DNA molecule under ~1 pN force and then reducing the force to ~0.2 pN. A rapid reduction in DNA extension would be observed if a single supercoilable DNA molecule was attached to the magnetic bead under observation. For the extension-rotation curves, we collected the DNA extension data at a rate of ~70 Hz for 20 seconds at every 2 turns between -40 and +40 turns.

Similar to previously published work(1), we calibrate the force applied to the DNA molecule by measuring the variation in horizontal motion of the magnetic bead for different magnet positions. We always start by measuring the extension-rotation curve for each DNA molecule in a 200 mM NaCl buffer. Next, the flow cells are washed with > 10* flow cell volume of the applicable co-solute buffer. After flowing in the new buffer, we always recalibrate the instrument to account for the change in refractive index due to presence of co-solute.

| Buffer | # of independent extension-force experiments (repeats) | # of independent extension-rotation experiments (repeats) |
| --- | --- | --- |
| 200 mM NaCl | 17 (2) | 14 (=>2) |
| 200 mM NaCl + 10% v/v Glycerol | 5 (2) | 5 (=>2) |
| 200 mM NaCl + 20% v/v Glycerol | 5 (2) | 5 (=>2) |
| 200 mM NaCl + 10% v/v Ethylene Glycol | 5 (2) | 3 (=>2) |
| 200 mM NaCl + 5% wt./v PEG 8000 | 4 (2) | 4 (=>2) |
| 200 mM NaCl + 10% wt./v PEG 8000 | 4 (2) | 4 (=>2) |
| 200 mM NaCl + 15% wt./v PEG 8000 | 3 (2) | 3 (=>2) |

**Table S1.** Details of independent experiments and repeats.

| Buffer | Slope of extension-rotation curve (nm/ΔLk) | | Twist persistence length of DNA (nm) | | Plectoneme-linking number stiffness (nm) | |
| --- | --- | --- | --- | --- | --- | --- |
|  | Positively supercoiled | Negatively supercoiled | From positively supercoiled | From negatively supercoiled | From positively supercoiled | From negatively supercoiled |
| 200 mM NaCl | 43 ± 1 | 14.5 ± 3 | 85.65 ± 3.7 | N/A | 20.2 ± 1.2 | N/A |
| + 10% v/v Glycerol | 41 ± 1 | 15 ± 1 | 69.7 ± 11.2 | N/A | 17.33 ± 0.8 | N/A |
| + 20% v/v Glycerol | 35 ± 2 | 9.5 ± 1 | 53.6 ± 6.7 | N/A | 13.7 ± 1.8 | N/A |
| + 10% v/v Ethylene Glycol | 41 ± 4 | 14.6 ± 2 | 86.5 ± 13 | N/A | 23.6 ± 2 | N/A |
| + 5% wt./v PEG 8000 | 37 ± 2 | 35 ± 3 | 94 ± 11.36 | 45.8 ± 6.2 | 17.3 ± 1.5 | 17.77 ± 3.8 |
| + 10% wt./v PEG 8000 | 40 ± 3 | 33 ± 1 | 80.65 ± 7.2 | 45.16 ± 7.2 | 20.1 ± 2.7 | 12.6 ± 1.3 |
| + 15% wt./v PEG 8000 | 34 ± 3 | 31 ± 3 | 76 ± 15 | 80.45 ± 11.1 | 16.5 ± 0.3 | 12.36 ± 2.15 |

**Table S2.** Calculated mechanical parameters required to model DNA supercoiling. Errors represent standard errors.

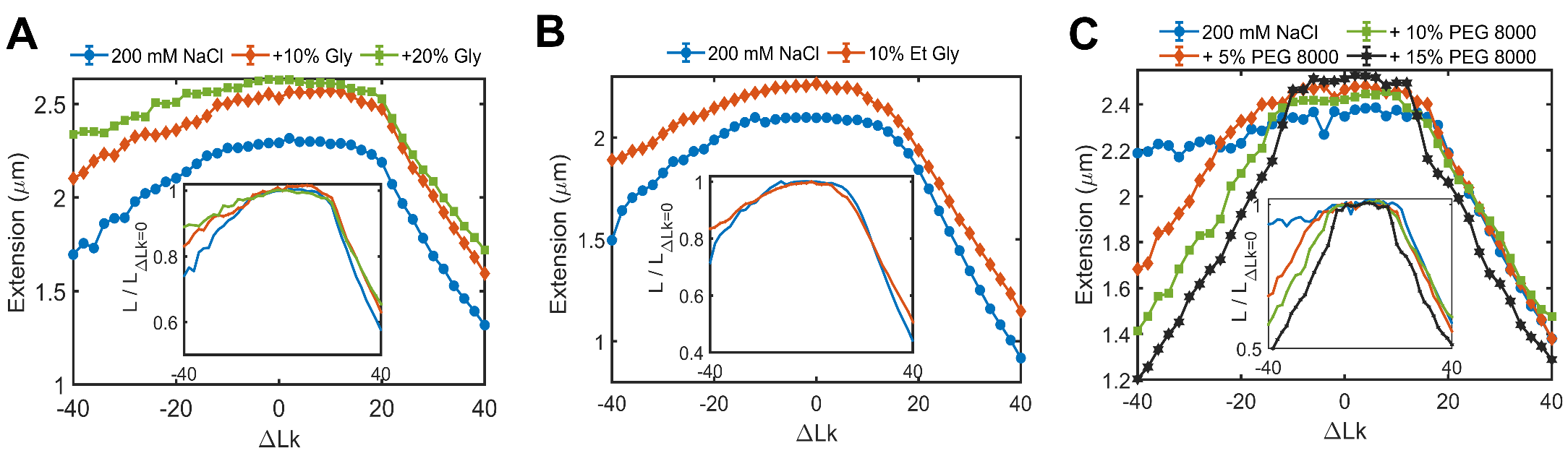
**S2. Representative hat curves in presence of co-solutes**

**FIG S1.** Representative extension-rotation curves demonstrate that presence of co-solutes affect DNA supercoiling. The extension-rotation curves presented in each sub-plot were collected on the same molecule. (A) Extension- rotation curve of DNA in 200 mM NaCl and 0% (blue), 10% (red) and 20% (green) v/v Glycerol (Gly) buffer. (B) Extension- rotation curve of DNA in 200 mM NaCl and 0% (blue) and 10% (red) v/v Ethylene Glycol (Et Gly) buffer. (C) Extension- rotation curve of DNA in 200 mM NaCl and 0% (blue), 5% (red), 10% (green) and 15% (black) wt./v Polyethylene Glycol 8000 (PEG) buffer. Inset show the extension-rotation curve normalized to the extension of a tortional relaxed DNA in the respective buffer conditions. Error bars represent standard errors and are smaller than the symbol size.

**S3. DNA bending persistence length and contour length in presence of co-solutes**

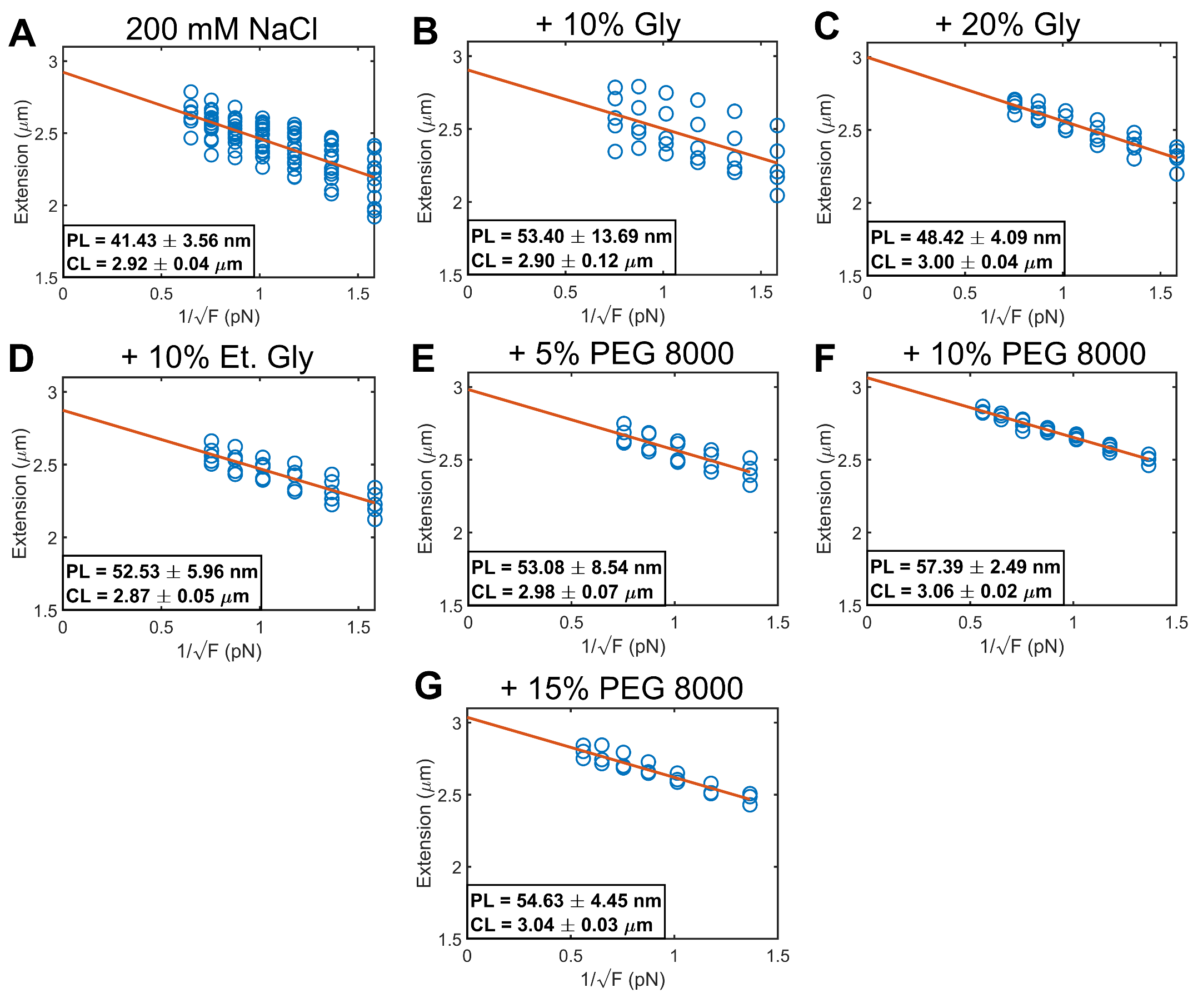

**Figure S2.** Persistence length (PL) and contour length (CL) of dsDNA in presence of different co-solutes. Persistence length was measured using the slope of the Extension- inverse of the square root of force curve(1). Extension (blue circles) was measure with measured at forces between ~0.8 pN and ~1.5 pN. Slope and intercept of the best linear fit line (red solid line) in plots of the DNA extension versus the inverse of the square root of force were used to determine the persistence length (PL), contour length (CL) and standard errors in accord with expectations from the worm-like chain polymer model.

**S4. Twist persistence length of DNA and twist stiffness of plectoneme**

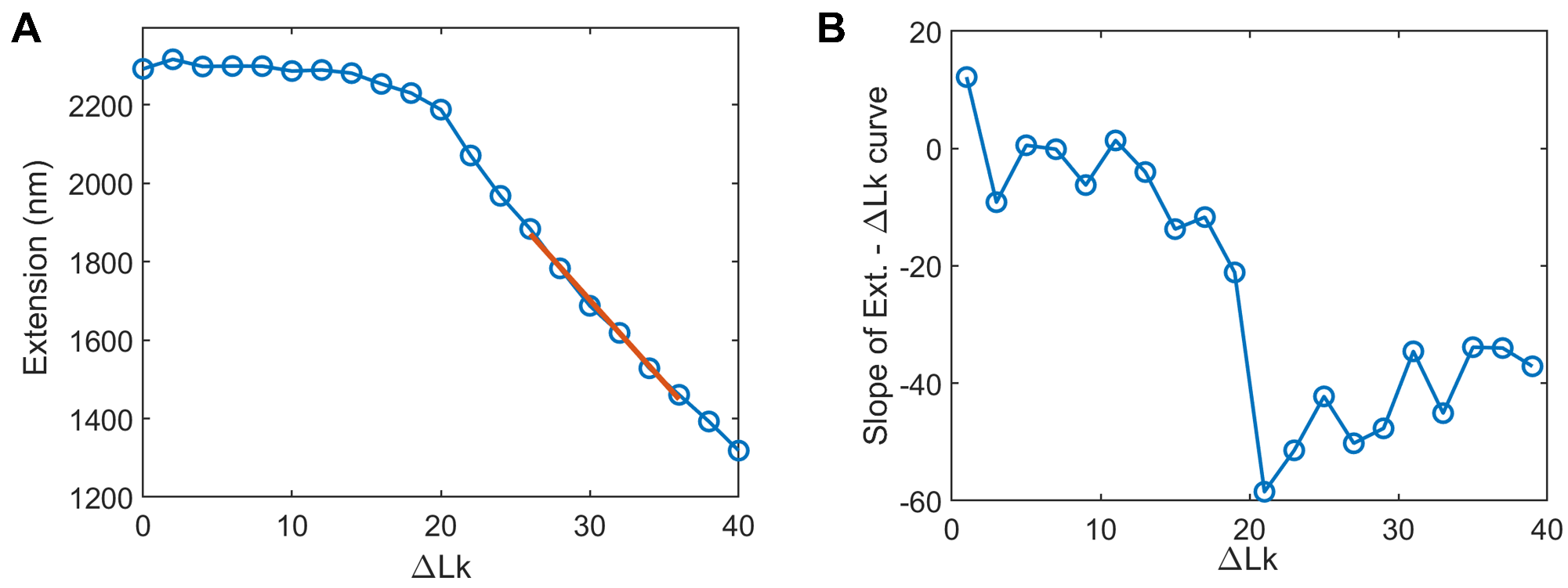

**Figure S3.** Extension-rotation curve is utilized to calculate the twist persistence length of DNA and plectoneme-linking number stiffness. (A) Representative extension-rotation curve of DNA under 0.8 pN of stretching force in 200 mM NaCl buffer. A straight line is fit to extension between ΔLk = 24 and ΔLk = 34. The line fit can be extrapolated to calculate the supercoiling density ($\sigma_{p}$) required for zero extension. (B) Slope of the extension-rotation curve. Supercoiling density at buckling ($\sigma_{s}$) can be calculated as minimum slope (first derivative) of the extension-rotation curve.

Quantitative modelling of DNA supercoiling can be performed using the bending persistence length of relaxed DNA ($A$), twist persistence length of DNA ($C$) and plectoneme-linking number stiffness ($P)$ (plectoneme-linking number stiffness is similar to the “twist stiffness of plectoneme” parameter defined in Marko et *al.*(2)). The bending persistence length of DNA is calculated as described in supplementary section S3. Model presented in Marko et *al.*(2) can be used to obtain the twist persistence length of DNA and twist stiffness of plectoneme from the extension-rotation curve. Marko et *al.*(2) describes the supercoiling density at buckling ($\left| \sigma_{s} \right|$) and supercoiling density for zero extension ($\left| \sigma_{p} \right|$) as

$$\left| \sigma_{s} \right| = \frac{1}{c_{s}} \left( \frac{2pg}{1-p/c_{s}} \right)$$

(S4.1)

$$\left| \sigma_{p} \right| = \frac{1}{p} \left( \frac{2pg}{1-p/c_{s}} \right)$$

(S4.2)

where, free energy of stretched and nicked DNA ($g$) under a stretching force ($f$) is $g = f - \left( \frac{k_{B}T}{A} \right)^{1/2}$. Twist stiffness of plectoneme $P = \frac{p}{k_{B}T{\omega_{0}}^{2}}$ & the contour-length rate of rotation of the relaxed double helix $\omega_{0}=\frac{2\pi}{helix repeat}$ .

The twist persistence length of DNA ($C$) is related to $c_{s}$ and defined as $c_{s}= k_{B}TC{\omega_{0}}^{2}- \frac{k_{B}TC^{2}{\omega_{0}}^{2}}{4A}\left( \frac{k_{B}T}{Af} \right)^{\frac{1}{2}}$. We obtain the buckling point ($\left| \sigma_{s} \right|$) by finding minimum of the first derivative of the extension-rotation curve (see supplementary Fig. S3 (B)). The model defines $\left| \sigma_{p} \right|$ as the supercoiling density when all of the DNA molecule is part of plectoneme and hence DNA extension is zero. To calculate $\left| \sigma_{p} \right|$, we first fit a straight line to the extension between ΔLk = 24 and ΔLk = 34. We then extrapolate the line to zero extension and calculate $\left| \sigma_{p} \right|$.

After calculating $\left| \sigma_{s} \right|$ and $\left| \sigma_{p} \right|$ from the extension-rotation curve, we can divide Eq. S4.1 by Eq. S4.2 to get

$$\frac{\left| \sigma_{s} \right|}{\left| \sigma_{p} \right|}= \frac{p}{c_{s}} or c_{s}= \frac{p \left| \sigma_{p} \right|}{\left| \sigma_{s} \right|}$$

(S4.3)

Eq. S4.3 can be used to simplify Eq. S4.1 to

$$p= \frac{2g}{\left| \sigma_{p} \right|^{2}-\left| \sigma_{p} \right|\left| \sigma_{s} \right|}$$

(S4.4)

Eq. S4.4 and Eq. S4.3 can be used to obtain $p$ and $c_{s}$, which can then be used to twist persistence length ($C$) and plectoneme-linking number stiffness ($P)$. The twist persistence length (~85 nm) and plectoneme-linking number stiffness (~20 nm) values of supercoiled DNA in 200 mM NaCl buffer are comparable to recent direct measurements by Gao et *al.* (3)

We only use $\left| \sigma_{s} \right|$ and $\left| \sigma_{p} \right|$ obtained from extension-rotation curve of positively supercoiled DNA because $\left| \sigma_{s} \right|$ is not well defined for negatively supercoiled DNA in the salt and force conditions used in the current paper.
